# Demonstration of chemotherapeutic mediated lymphatic changes in meningeal lymphatics in vitro, ex vivo, and in vivo

**DOI:** 10.1101/2024.01.06.574460

**Authors:** L. Monet Roberts, Jennifer H Hammel, Francesca Azar, Tzu-Yu (Alkaid) Feng, Jessica J. Cunningham, Melanie Rutkowski, Jennifer Munson

**Affiliations:** Fralin Biomedical Research Institute, Roanoke, VA 24016, USA; Department of Biomedical Engineering and Mechanics, Virginia Tech, Blacksburg, VA 24061, USA; Department of Microbiology, Immunology, and Cancer Biology, School of Medicine, University of Virginia, Charlottesville, VA 22903, USA

**Author notes:** **Contact information:** Jennifer Munson, Associate Professor, Fralin Biomedical Research Institute, Virginia Tech, Room 1210, 4 Riverside Circle, Roanoke, VA 24016. Authors contributed equally (L. Monet Roberts, Jennifer H Hammel).

## Abstract

Systemic chemotherapeutics target cancer cells but are also known to impact other cells away from the tumor. Questions remain whether systemic chemotherapy crosses the blood-brain barrier and causes inflammation in the periphery that impacts the central nervous system (CNS) downstream. The meningeal lymphatics are a critical component that drain cerebrospinal fluid from the CNS to the cervical lymph nodes for immunosurveillence. To develop new tools for understanding chemotherapy-mediated effects on the meningeal lymphatics, we present two novel models that examine cellular and tissue level changes. Our in vitro tissue engineered model of a meningeal lymphatic vessel lumen, using a simple tissue culture insert system with both lymphatic endothelial and meningeal cells, examines cell disruption. Our ex vivo model culturing mouse meningeal layers probes structural changes and remodeling, correlating to an explant tissue level. To gain a holistic understanding, we compare our in vitro and ex vivo models to in vivo studies for validation and a three-tier methodology for examining the chemotherapeutic response of the meningeal lymphatics. We have demonstrated that the meningeal lymphatics can be disrupted by systemic chemotherapy but show differential responses to platinum and taxane chemotherapies, emphasizing the need for further study of off-target impacts in the CNS.

## Introduction

The meningeal lymphatics, which drain the central nervous system (CNS) ^1,2^, have been implicated in a variety of CNS diseases. The meninges in which they reside consist of three protective layers surrounding the brain that provide insulation between the skull and the brain, serve as an immune cell reservoir, and drain cerebrospinal fluid (CSF) from the CNS. Within the meninges, the lymphatic system consists of dural and basal meningeal lymphatic vessels that regulate immune cell trafficking and allow CSF drainage to the cervical lymph nodes. Previous studies show that the meningeal lymphatics are involved in brain tumor drainage ^3,4^, Alzheimer’s disease ^5,6^, Parkinson’s disease, and traumatic brain injury ^7^ with conservation between humans and mice ^8^. Further, the meningeal lymphatics are critical in regulating immune response following radiotherapy ^9^, and vascular endothelial growth factor receptor (VEGFR) inhibition through sunitinib treatment leads to regression in meningeal lymphatic vessels ^10^. Though there are implications in cancer and cancer-related sequelae, the degree to which the meningeal lymphatics contribute to CNS pathophysiology in response to chemotherapeutic treatment has yet to be determined.

Chemotherapy is one of the most conventional forms of cancer treatment as a primary or adjuvant therapy following resection and/or radiotherapy. It has increased survival rates since its inception for a number of cancers globally. However, there are still known and unknown risks and side effects in response to chemotherapeutics ^11,12^. Most chemotherapeutics are intravenously injected into patients. They circulate throughout the body and enter the tumor through the blood vasculature but come into contact with all tissues. After entering the tissue, chemotherapy drains via the lymphatic system, where it can have counter-therapeutic effects. Systemic chemotherapy recapitulating a breast cancer treatment regimen using either carboplatin or docetaxel in pre-clinical models leads to lymphangiogenesis in the primary tumor site and draining lymph nodes that can promote invasion of cancer cells^11,12^. However, the potential impact of chemotherapy on the distant meningeal lymphatics following systemic circulation in breast cancer is not fully defined.

With the renewed interest in the meningeal lymphatics, the development of in vitro models is crucial to answering biological questions and understanding the physiology of this essential lymphatic system. In vitro models allow the examination of crosstalk between cell types in an isolated and customizable system ^13,14^. Here we have developed an in vitro human cell–based model of the meningeal lymphatics and an ex vivo culture of mouse meningeal layers and compared these models to in vivo studies. These models can provide future insights to better inform treatment of patients and efforts to ameliorate cognitive deficits post-treatment and increase patient quality of life.

## Materials and Methods

### Cell lines

4T1 murine mammary carcinoma cells were purchased from the ATCC. Cell lines were authenticated using MAP testing and bi-monthly mycoplasma testing to ensure they were free of viral and other pathogens. To limit genetic drift, cells were maintained at low passage numbers and maintained as frozen stocks at −180 °C and expanded only for inoculation into mice. Tumor cell lines were cultured according to conditions outlined by ATCC. Human lymph node lymphatic endothelial cells (hLECs) (Sciencell) were cultured on fibronectin coated flasks in VascuLife® VEGF-Mv media (Lifeline Cell Technology). Human meningeal cells (hMCs) were cultured on poly-L-lysine coated flasks in MenCM complete media (Sciencell).

### Animals

All animals were maintained in specific pathogen-free barrier conditions at the University of Virginia. All procedures complied with regulations of the Institutional Animal Care and Use Committee at Virginia Polytechnic Institute & State University and University of Virginia. Female BALB/c mice, C57BL/6, and Prox1-Tdtomato mice were purchased from Jackson Laboratories. Mice tumor implantation and chemotherapeutic treatments were performed as previously described ^15,16^. Briefly, BALB/c mice were injected with 10^4^ 4T1 murine mammary carcinoma cells. Tumor cells were resuspended in a mixture of ice-cold phosphate buffered saline (PBS) and 3.3 mg/ml Matrigel, in a total volume of 50 μl, into the fourth mammary fat pad. Tumor growth was monitored, and when tumors reached a volume of 10 mm total dimension as measured using calipers, mice were monitored daily. If tumors reached no more than 15 mm total diameter, became ulcerated, or impaired the mouse’s ability to ambulate, eat or drink, it was euthanized. Otherwise, all analyses were performed 20-25 days post tumor injection. Beginning on day 7, carboplatin and docetaxel were administered intravenously using tail-vein injection at 8mg/kg of body weight with mice administered saline/20% ethanol as a control every other day over the course of 6 days.

### Tissue processing, harvesting, imaging, and analysis of in vivo and ex vivo meningeal layers

For in vivo studies, mice were euthanized via carbon dioxide asphyxiation and secondary euthanasia through intracardiac perfusion with PBS and subsequent intracardiac perfusion with 4% paraformaldehyde (PFA). Skull caps were collected and incubated for 18 h at 4°C in 4% PFA. Following 18 h incubation, skull caps were transferred to fresh PBS and stored at 4°C until harvesting of meningeal layers. For ex vivo tissues, dissected skull cap explants were transferred directly to explant media (DMEM + L-Glutamax (10567-014; Fisher Scientific), 10% FetalPURE (Genesee Scientific, 2.5% penicillin-streptomycin and 2.5% 250 ug/mL amphotericin B ^17^ on ice until subsequent chemotherapeutic treatments of either DMSO or 1 μM of carboplatin (Tocris Bioscience), 1 μM docetaxel (Tocris Bioscience), or in combination in explant media and incubated at 37°C for 24 h. Following chemotherapeutic treatment, skull caps were fixed for 18 h in 4% PFA at 4°C. Following fixation, skull caps were rinsed with PBS and stored in fresh PBS at 4°C. Harvesting of dural whole-mount meningeal layers was achieved through dissection from skull caps. The meningeal layers were dissected from the occipital side of the skull cap, peeled from the skull, and stored in PBS at 4°C until subsequent processing for immunostaining.

For live whole mount ex vivo meningeal layer dissection and labeling for imaging, meningeal layers were harvested from Prox1-Tdtomato mice without perfusion and gently dissected from skull caps using a Zeiss Axio Zoom microscope in sterile explant media with 5X Anti/Anti (Thermo). The layers were transferred to a poly-L-lysine coated 24 well glass bottom dish with sterile Ringer’s solution (102 mM NaCl, 5 mM KCl, 2 mM CaCl2, 28 mM sodium lactate) with 1X Anti/Anti and flattened out as much as possible. 2 µM FITC docetaxel (Conju-Probe) or DMSO in sterile Ringer’s/Anti/Anti solution was added to the well plate at equal volume to get a final concentration of 1 µM with Click-iT™ EdU 647 solution per protocol (ThermoFisher). Time-lapse images were acquired in 15 min intervals over 6 h before remaining 12 h incubation before fixation and further immunofluorescence staining and EdU detection as described below.

### Tissue culture insert model

12-mm tissue culture inserts with 8 µm pore size (Corning) were inverted in 6 well plates. LECs were seeded on the underside of the insert at 50,000 per insert and cultured for 2 h before re-inverting in a tissue culture plate. After 48 h, 75,000 hMCs were seeded within the tissue culture insert. The co-culture model was grown for an additional 24 h to reach confluence before assays were performed. At confluence, models were treated with 1 µM carboplatin, 1 µM docetaxel, a combination of both, or vehicle (DMSO) in Vasculife® for 24 h.

### Permeability assays

300 µL of 10 µg/mL FITC dextran 10,000 MW, anionic, lysine fixable (Invitrogen) in basal Vasculife® was added inside the tissue culture insert. 300 µL of basal Vasculife® was placed underneath the insert. After 6 h, media from underneath the insert was collected and analyzed on a CLARIOstar plate reader.

### Immunofluorescence

Tissue culture inserts were fixed in 4% PFA for 15 min. The membranes of each insert were carefully cut out and mounted on positively charged slides for staining. Donkey serum was used to block nonspecific binding. Membranes were stained with sheep anti-human CD31 (R&D) at a 1:40 dilution and mouse anti-human alpha smooth muscle actin (Abcam) at a 1:200 dilution for 2 h at room temperature (RT) followed by donkey anti-sheep Alexa Fluor 647 secondary antibody (Invitrogen) at a 1:400 dilution and donkey anti-mouse Alexa Fluor 555 antibody (Invitrogen) at a 1:1000 dilution for 1 h at RT. Membranes were then stained with 488 Phalloidin for F-actin (Invitrogen) at a 1:400 dilution and DAPI (Invitrogen) at a 1:5000 dilution for 45 min. Slides were then mounted with Fluoromount, allowed to dry overnight, sealed with nail polish, and stored at 4°C. Slides were imaged on a Zeiss Axio Observer.

For meningeal whole mounts, tissues were stained in 24 well plates until mounting. Specifically, tissues were blocked in blocking buffer solution containing 0.5% triton-X 100, 2% bovine serum albumin, and 3% donkey serum in PBS for 1-2 h at RT. Tissues were then incubated in the following primary antibodies: LYVE-1 (Abcam) at a dilution of 1:100, Prox1 (Abcam) at a dilution of 1:200, α-SMA (Abcam), at a dilution of 1:100, and CD31 (R&D Systems) at a dilution of 1:100 at 4°C overnight. Tissues were rinsed 2X for 5 min each in permeabilization buffer and then incubated in donkey anti-mouse FITC (Fisher Scientific), donkey anti-rabbit Alexa Fluor 555 (Invitrogen), and donkey anti-goat Alexa Fluor 647 (Invitrogen) at dilutions of 1:200 for 1 h at RT. Tissues were then rinsed 2X in permeabilization solution and 2X in PBS. Tissues were incubated with DAPI for 5 min and then rinsed in PBS 2X for 10 min each and mounted on glass slides and allowed to dry completely. Images were acquired at 10X magnification using the Zeiss Axio Observer. At least five regions of interest were imaged across the superior sagittal sinus and along the transverse sinus.

### Proliferation assay

The Click-iT™ EdU Cell Proliferation Kit for Imaging, Alexa Fluor™ 488 dye (ThermoFisher) was utilized to quantify proliferation in the meningeal lymphatic (ML) model. During the last 6 h of culture, 10 µM EdU was added to the culture media. At 24 h, tissue culture inserts were fixed in 4% PFA. EdU was then detected as specified by the manufacturer. Finally, insert membranes were stained for CD31 and αSMA as described above.

For EdU detection in the meningeal layer whole mounts, the Click-iT™ EdU Cell Proliferation Kit for Imaging, Alexa Fluor™ 647 dye (ThermoFisher) for timelapse imaging was used for 6 h of culture sterile in Ringer’s solution/1X Anti/Anti solution and removed and replaced with sterile Ringer’s solution/1X Anti/Anti solution for remaining 12 h. After 18 h, the meningeal layers were fixed with 4% PFA and underwent subsequent immunofluorescence staining with LYVE-1 and DAPI as described above. EdU was then detected as specified by the manufacturer.

### Image Quantification

For total area of fluorescence in the tissue engineered model, images were thresholded using Fiji and total area of signal was measured. For proliferation, total nuclei were quantified via DAPI stain and proliferating cells were quantified via positive EdU staining. Proliferation was reported as a percentage of EdU+ cells out of total cell number. Analysis of lymphatic vessels was performed in the dural sinuses, specifically the superior sagittal sinus and transverse sinus, respectively. Branching points and intussusceptions/loops were hand-counted across the vessels with total or average branching points and average intussusceptions being reported. Diameter measurements of the meningeal vessels were calculated through a CellProfiler pipeline ^18^ trained to identify vessels as objects. To determine vessel diameter, the ‘meanRadius’ measurement for each vessel object from the ‘MeasureObjectSizeShape’ module in CellProfiler was multiplied by two.

### Statistical analysis

One-way ANOVA or two-way ANOVA was performed for groups of two or more with statistical significance indicated by *p* < 0.05. When significance was identified, ad hoc *t-*tests (ratio paired or unpaired) were performed to assess significance between groups with *p* < 0.05 indicating significance. All graphs represent mean ± standard error mean. Individual statistical results are discussed with the presented data.

### Data availability

Data presented in the study are included in the article and supplementary material. Data and CellProfiler pipeline are available upon request and further inquiries can be directed to the corresponding author.

## Results

### Systemic chemotherapy induces morphological changes in lymphatic vasculature in vivo and ex vivo

4T1 tumor-bearing mice were treated with systemic 1 µM platinum or taxane chemotherapy or DMSO as the vehicle prior to dural meningeal harvest (Figure 1). Chemotherapy-induced changes in meningeal layers were observed through lymphatic vessel remodeling regarding diameter, intussusceptions/loops, and branching points (Figure 2A) in the dural sinuses known as the superior sagittal sinus (SSS) and the transverse sinus (Supplementary Video S1, Supplementary Figure 1 & 2, Figure 2A). Specifically, intussusceptions/loops and diameter of the lymphatic vessels were decreased in the presence of taxane chemotherapy compared to both vehicle and platinum therapy (Figure 2B-D, Supplementary Figure S3A-C). Similarly, branch remodeling revealed a lower trend in presence of taxane chemotherapy (Figure 2D).

**Figure 1.**
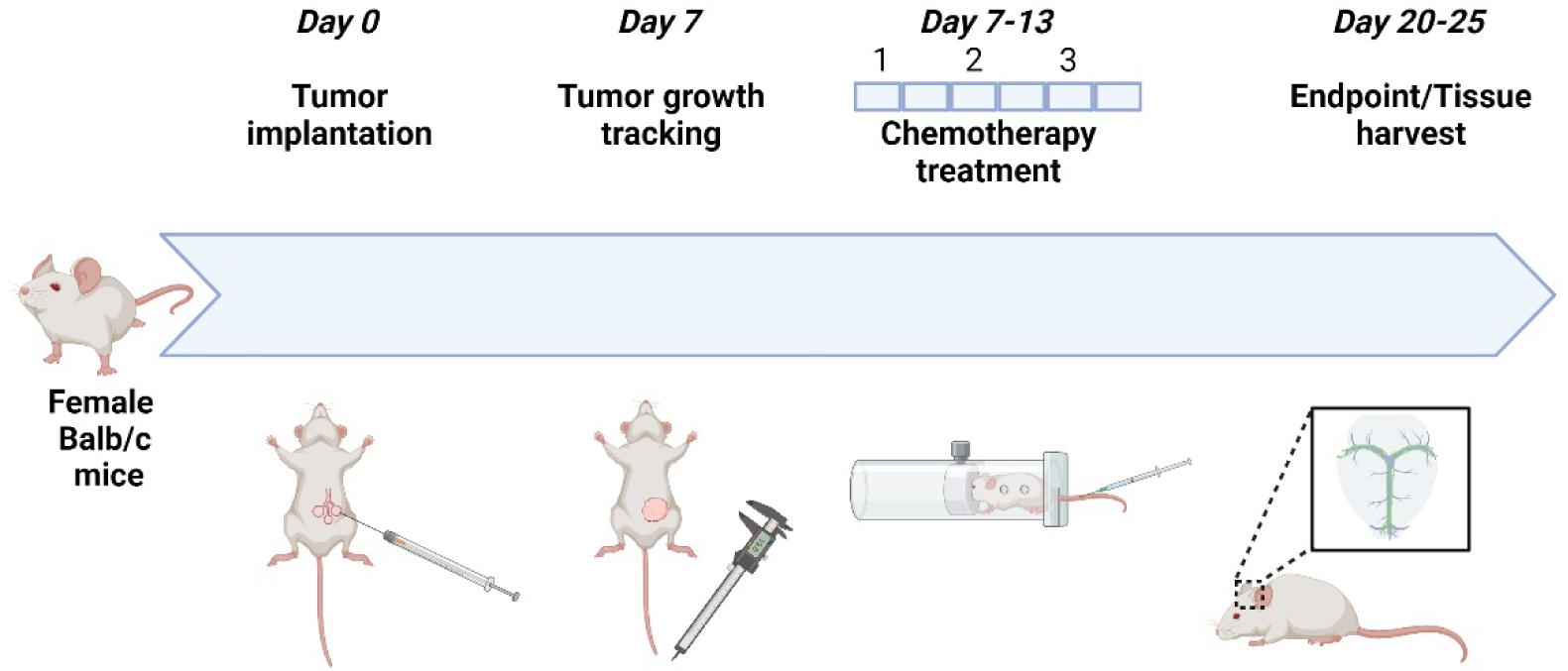
Implantation, treatment, and meningeal layer harvest from mice and remodeling outcomes. Schematic representation of timeline of tumor implantation, tumor growth tracking with a caliper, chemotherapeutic treatment through tail-vein injection, and tissue harvesting.

**Figure 2.**
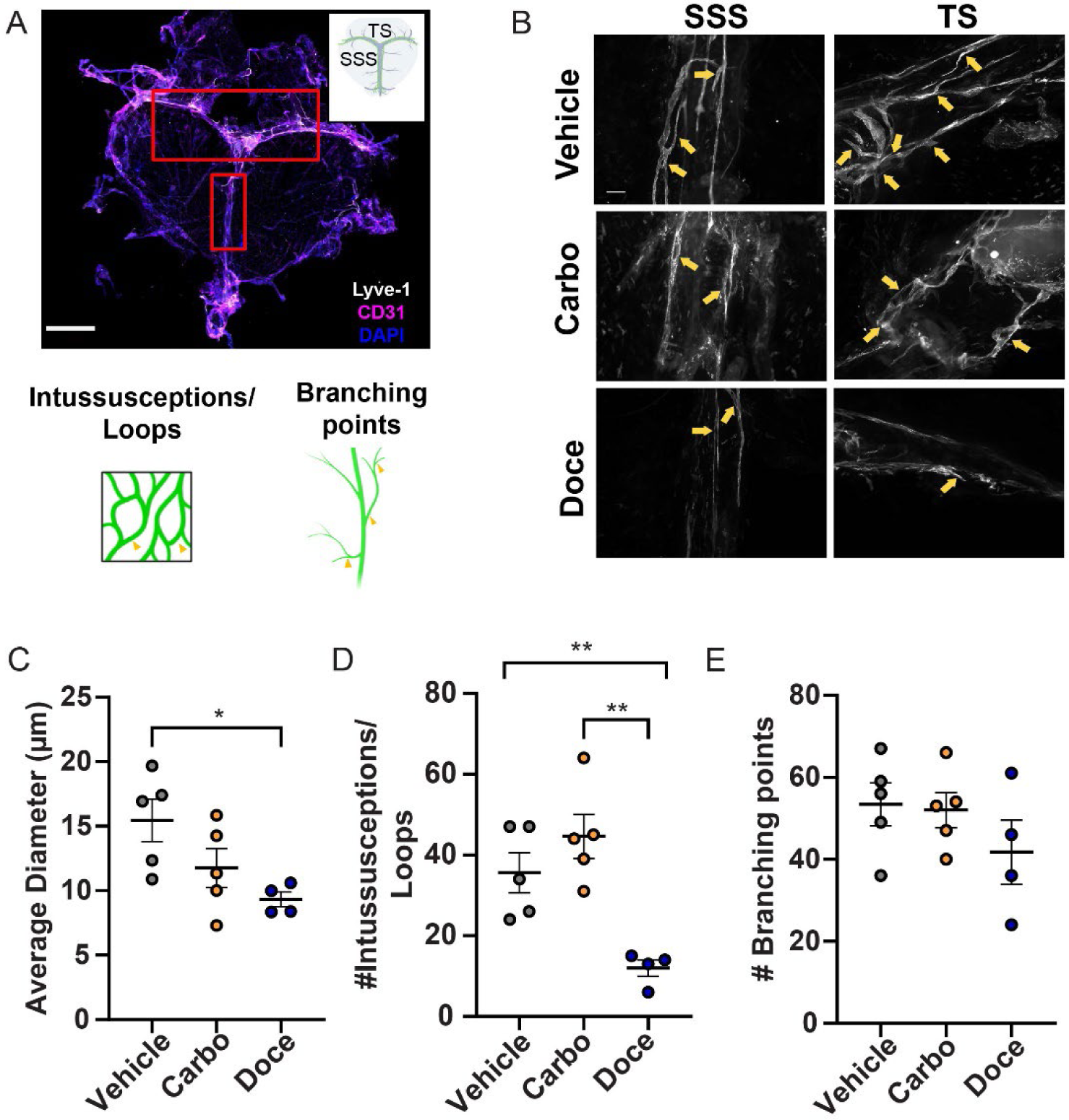
Meningeal lymphatic remodeling varies in response to different chemotherapeutic agents. (A) Representative image of a meningeal layer highlighting the superior sagittal sinus (SSS) and transverse sinus (TS) (red boxes; inset is cartoon representation of meningeal layer with the SSS and the TS denoted) (scale bar: 2 mm). Schematic representing meningeal lymphatic vessel remodeling outcomes for analysis: intussusceptions/loops (left) and branching (right) (features denoted by yellow arrows). (B) Representative images of remodeling outcomes of intussusceptions/loops and branching points in meningeal layers in the presence of systemic vehicle, carboplatin, and docetaxel (features denoted by yellow arrows) (scale bar: 100 µm). (C) Quantification of average diameter of meningeal lymphatic vessels in the presence of chemotherapies and vehicle. (D) Quantification of intussusceptions/loops and (E) branching in meningeal layers. All graphs represent Mean ± SEM (n =5 for vehicle and carboplatin, n =4 for docetaxel; *, p < 0.05, **, p < 0.01)

The meningeal vessels residing in the SSS and the TS are initiating vessels that receive interstitial fluid ^19^. We wanted to explore if there were differences in remodeling between the SSS and the TS. There was no difference in the diameter of vessels or branching within the SSS and the TS (Supplementary Figure S3B,C). We observed a significant decrease in intussusceptions/loops in the presence of docetaxel compared to carboplatin and vehicle in the TS (Supplementary Fig S3C). We then wanted to validate whether direct chemotherapeutic treatment would have an effect on meningeal lymphatic remodeling. We harvested murine meningeal layers and then cultured ex vivo in the presence of the vehicle or 1 µM of carboplatin, docetaxel, or vehicle (Figure 3A). Branching and intussusceptions were significantly decreased in docetaxel-treated meningeal layers compared to vehicle and carboplatin culture in ex vivo mouse meningeal lymphatic layers (Figure 3B-D) similar to what we observed in vivo. Overall, systemic docetaxel was deleterious to the dural meningeal lymphatic vessel remodeling.

**Figure 3.**
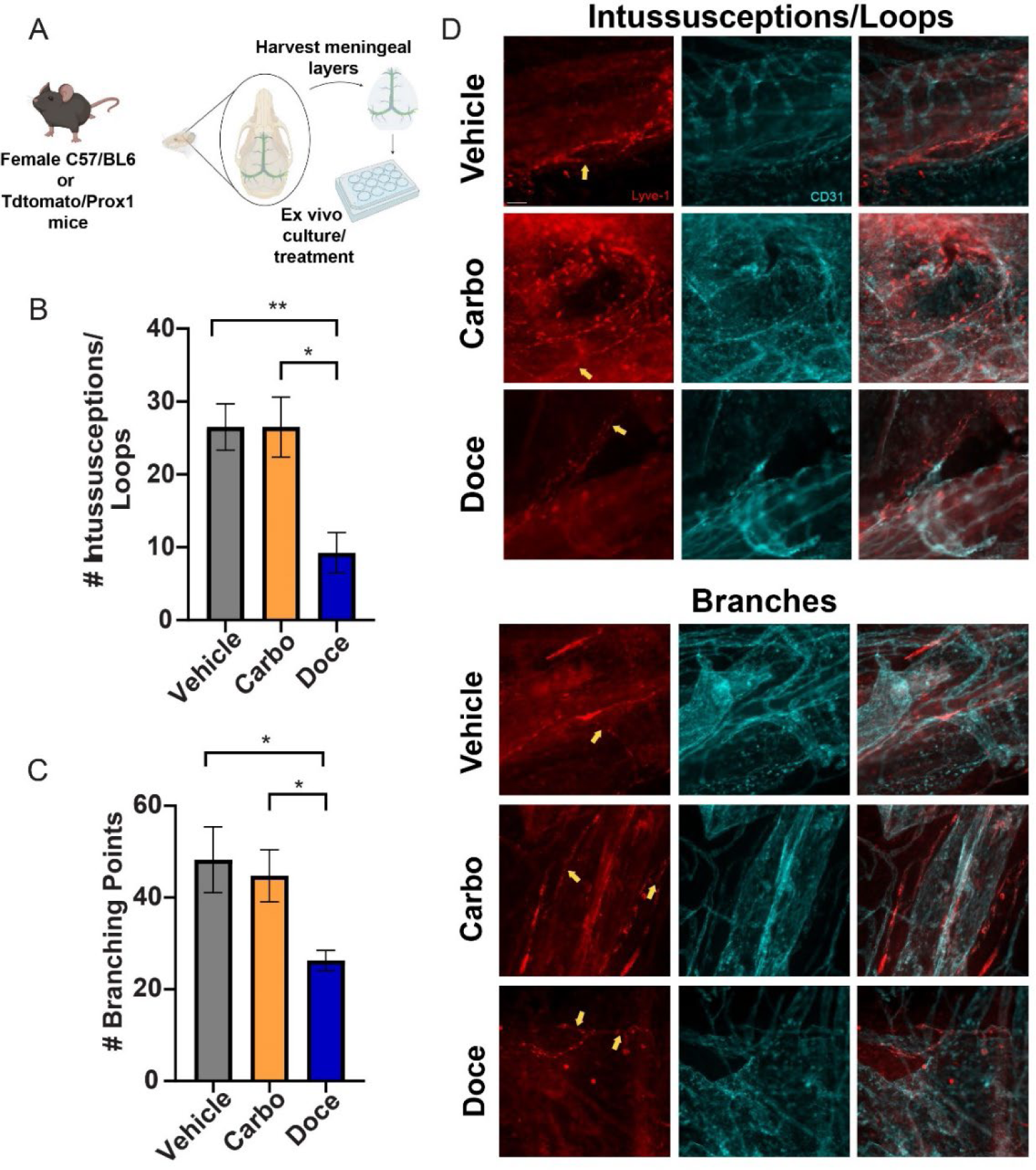
Docetaxel decreases lymphatic remodeling in ex vivo murine meningeal layers. (A) Representative images of intussusceptions/loops in ex vivo meningeal layers in the presence of 1 µM carboplatin, 1 µM docetaxel, or vehicle. (B) Quantification of intussusceptions/loops in meningeal layers in the presence of carboplatin, docetaxel, or vehicle (n=4). (C) Quantification of branching points in meningeal layers in the presence of carboplatin, docetaxel, or vehicle (n=4). (D) Representative images of branching points in meningeal layers in the presence of carboplatin, docetaxel, or vehicle. All graphs represent Mean ± SEM, *, p < 0.05, **, p < 0.01)

### Development of in vitro meningeal lymphatics model

To represent the lumen of a meningeal lymphatic model, LECs are seeded on the underside of a tissue culture insert. Then, the top of the insert is seeded with meningeal cells (Figure 4A). Initial work aimed to optimize a co-culture media for the model for ease of future applications with more complexity. Medias tested include AIM-V with VEGF-C and EGF supplementation and Neurobasal media with Glutamax, B27, and N2 (Gibco). Preliminary data showed LECs had decreased viability in all tested media compared to Vasculife (Supplementary Figure S4). However, meningeal cells were viable in LEC media (Figure 4B) and had significantly more live cells per FOV (Figure 4C); thus, this media was selected for co-culture. The barrier function of the model compared to monoculture controls was examined via dextran permeability assay, demonstrating that meningeal fibroblasts have higher barrier function than lymph node (LN) LECs (Figure 4D). Distinct monolayers are visualized in the complete model via CD31 and F-actin staining, demonstrating menial cell transmigration (Figure 4E).

**Fig 4.**
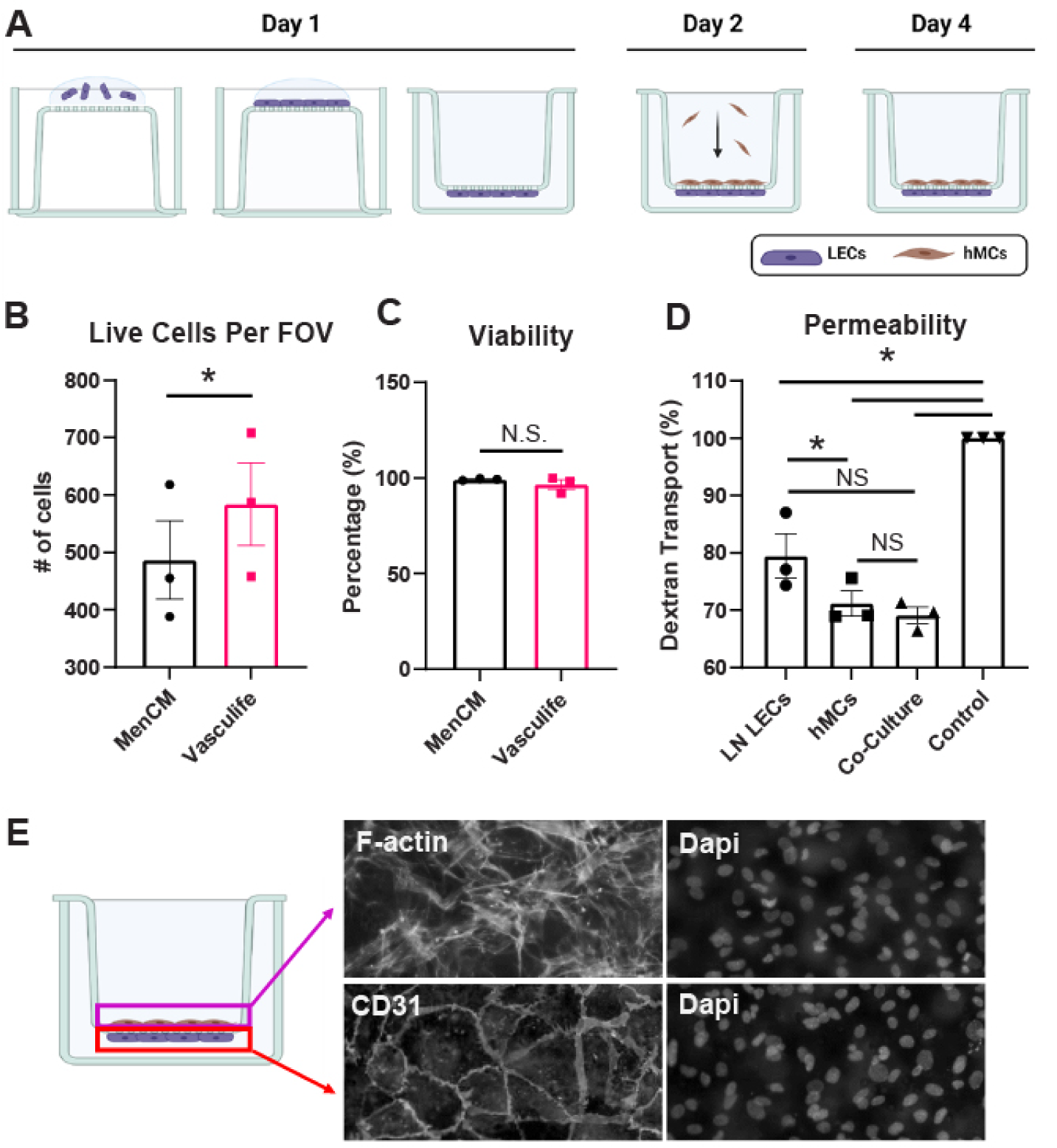
In vitro meningeal lymphatic model development. LN LECs are seeded on the underside of a tissue culture insert for 2 h before inversion and culture for 24 h. Then, meningeal cells are added into the tissue culture insert and cultured an additional 36 h. (A). Suitability of co-culture media is verified via total meningeal cells per field of view (B) and viability of meningeal cells in Vasculife (C). Permeability is quantified via dextran transport assay (D). Representative images of each monolayer are shown, with meningeal cells stained for F-actin (magenta) and LECs stained for CD31 (red). Nuclei, stained with DAPI, are shown in cyan (E). Data shown are biological replicates (n=3), mean +/− SEM, with * denoting p<0.05. Scale bar is 50 µm.

### LEC-MC crosstalk alters response to platinum and taxane chemotherapy treatment

The meningeal lymphatics model was then utilized to examine response to carboplatin and docetaxel treatment. Representative images demonstrate that MCs are highly aligned and elongated when in a confluent monolayer under vehicle and carboplatin treatment (Figure 5A). Alone, MCs demonstrate the beginnings of alignment (Supplementary Figure S5A). However, docetaxel treatment causes morphological changes, with the MCs beginning to form clumps or plaques (Figure 5A, Supplementary Figure S5A). When quantified, MCs formed a significantly more complete monolayer when in co-culture than when cultured alone (Figure 5B). This effect was also seen during chemotherapy treatment (Figure 5B). Carboplatin did not significantly decrease the MC coverage compared to the vehicle but showed a more complete monolayer in co-culture than alone (Figure 5B). MCs also had higher MC coverage in co-culture than alone when treated with docetaxel (Figure 5B). Further, docetaxel caused significant disruption of the monolayer compared to the vehicle and carboplatin, for both co-culture and MCs alone (Figure 5B). Additionally, MC count was not significantly different when alone vs in co-culture in the vehicle or carboplatin controls but was higher in co-culture for docetaxel treated groups (Supplementary S5B), suggesting differences in cell spread. Overall, co-culture with LECs had a positive impact on MC coverage and showed decreased disruption during chemotherapy treatment.

**Fig 5.**
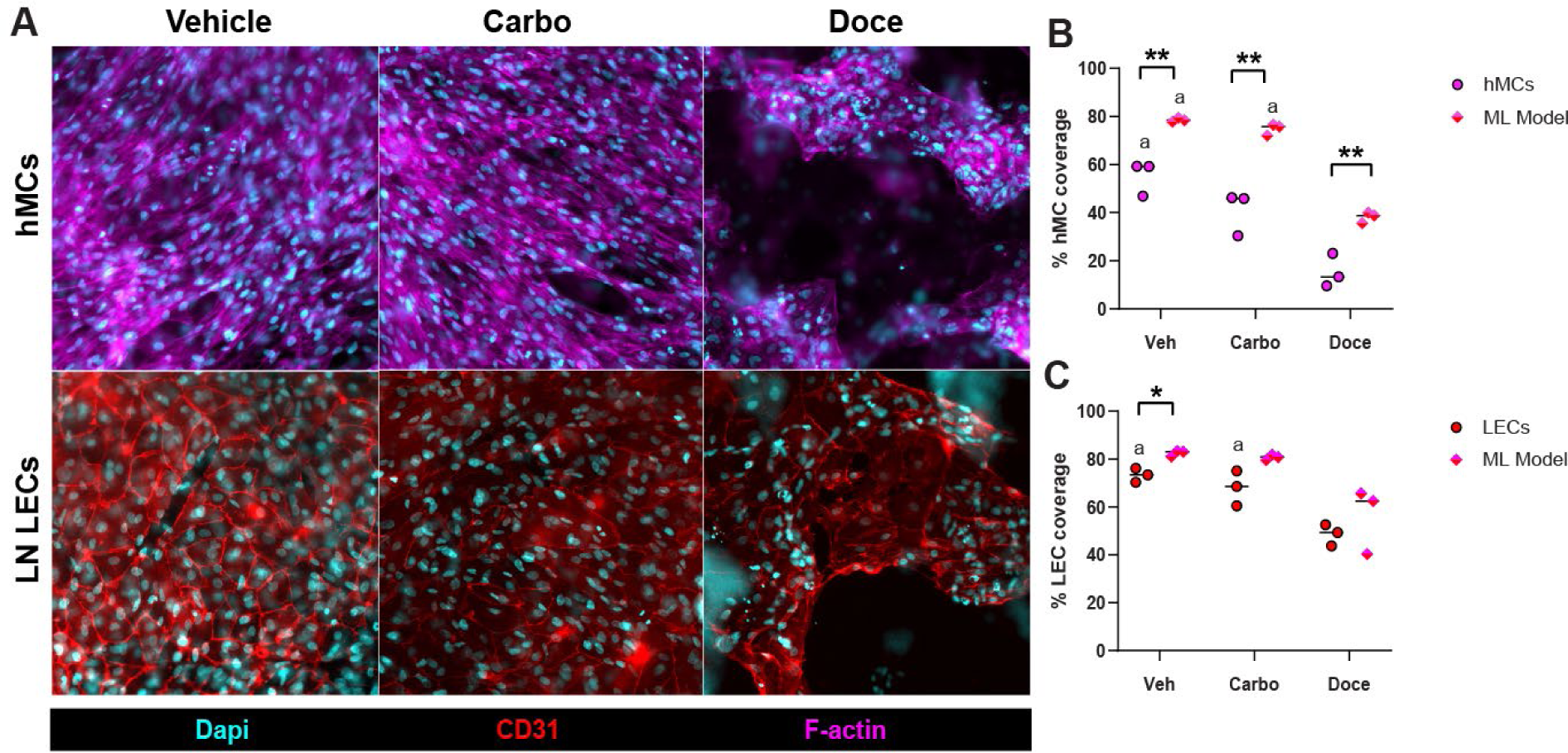
Docetaxel disrupts LEC and hMC coverage. ML models are treated with 1 µM carboplatin, 1 µM docetaxel, or vehicle (DMSO) for 24 hours. Representative images of ML models are shown, with nuclei stained with DAPI (cyan), hMCs stained for F-actin (magenta) and LECs stained for CD31 (red). Scale bar is 50 µm (A). Coverage of hMcs (B) and LECs (C) is quantified in ImageJ and reported as percentage, within the ML model and as monocultures. Data shown are biological replicates (n=3) and mean, with * denoting p<0.05, ** denoting p<0.01. a denotes significant difference from docetaxel treatment.

LECs showed less benefit from co-culture during chemotherapy treatment. Representative images demonstrate similarly intact monolayers of LECs (Figure 5A, Supplementary Figure S5A) in vehicle and carboplatin groups. Docetaxel treated groups show incomplete LEC monolayers and altered CD31 staining (Figure 5A, Supplementary Figure S5A). In the vehicle group, LEC coverage was significantly higher in co-culture than when cultured alone (Figure 5C). However, LEC coverage (Figure 5C) and LEC count (Supplementary Figure S5C) was not significantly different between co-culture and the LEC only control for any of the chemotherapy groups. Carboplatin was shown not to disrupt LEC monolayer coverage compared to vehicle controls (Figure 5C). Docetaxel significantly decreased LEC coverage in co-culture and alone compared to the vehicle (Figure 5C). Compared to carboplatin, docetaxel only significantly decreased LEC coverage in LECs cultured alone (Figure 5C).

In addition to coverage, LEC junctions, area, and aspect ratio were examined. LECs had more disrupted junctions when cultured alone than in co-culture in the vehicle and docetaxel groups (Supplementary Figure S6A,B). Chemotherapy did not increase junction disruption in LEC monocultures (Supplementary Figure S6A,B). However, in co-culture, LECs had more disrupted junctions when treated with both carboplatin and docetaxel (Supplementary Figure S6A,B). LECs were shown to be more elongated when cultured alone compared to co-culture when treated with carboplatin (Supplementary Figure S6C). In the meningeal lymphatics model, LECs were more elongated when treated with docetaxel compared to the vehicle (Supplementary Figure S6C). LECs had significantly higher area in the LEC only control than co-culture (Supplementary Figure S6D). These data further suggest differences in cell spread and size during co-culture and chemotherapy treatment.

### Total proliferation is altered in vitro, spatial distribution of proliferation altered ex vivo

Proliferation was also altered by crosstalk and chemotherapy. MCs trended toward higher proliferation alone than in co-culture in the vehicle control, but this effect was not significant (Figure 6A,B, Supplemental Figure S7). With carboplatin treatment, proliferation was significantly lower in MCs in co-culture compared to MCs cultured alone (Figure 6A,B, Supplemental Figure S7). Docetaxel-treated groups had very low levels of proliferation in co-culture and alone and were significantly lower than vehicle and carboplatin groups (Figure 6A,B, Supplemental Figure S7). LECs were shown to have higher proliferation in co-culture than alone for the vehicle (Figure 6A,C, Supplemental Figure S7). Carboplatin treatment showed the same trend of higher proliferation in co-culture than alone (Figure 6A,C, Supplemental Figure S7). However, there was no significant difference in LEC proliferation between co-culture and LECs alone for docetaxel (Figure 6A,C, Supplemental Figure S7). Interestingly, proliferation in the LEC-only control was not significantly altered by any chemotherapy treatment (Figure 6C, Supplemental Figure S7). However, LEC proliferation in co-culture was significantly lower in docetaxel compared to both vehicle and carboplatin (Figure 6A,C).

**Figure 6.**
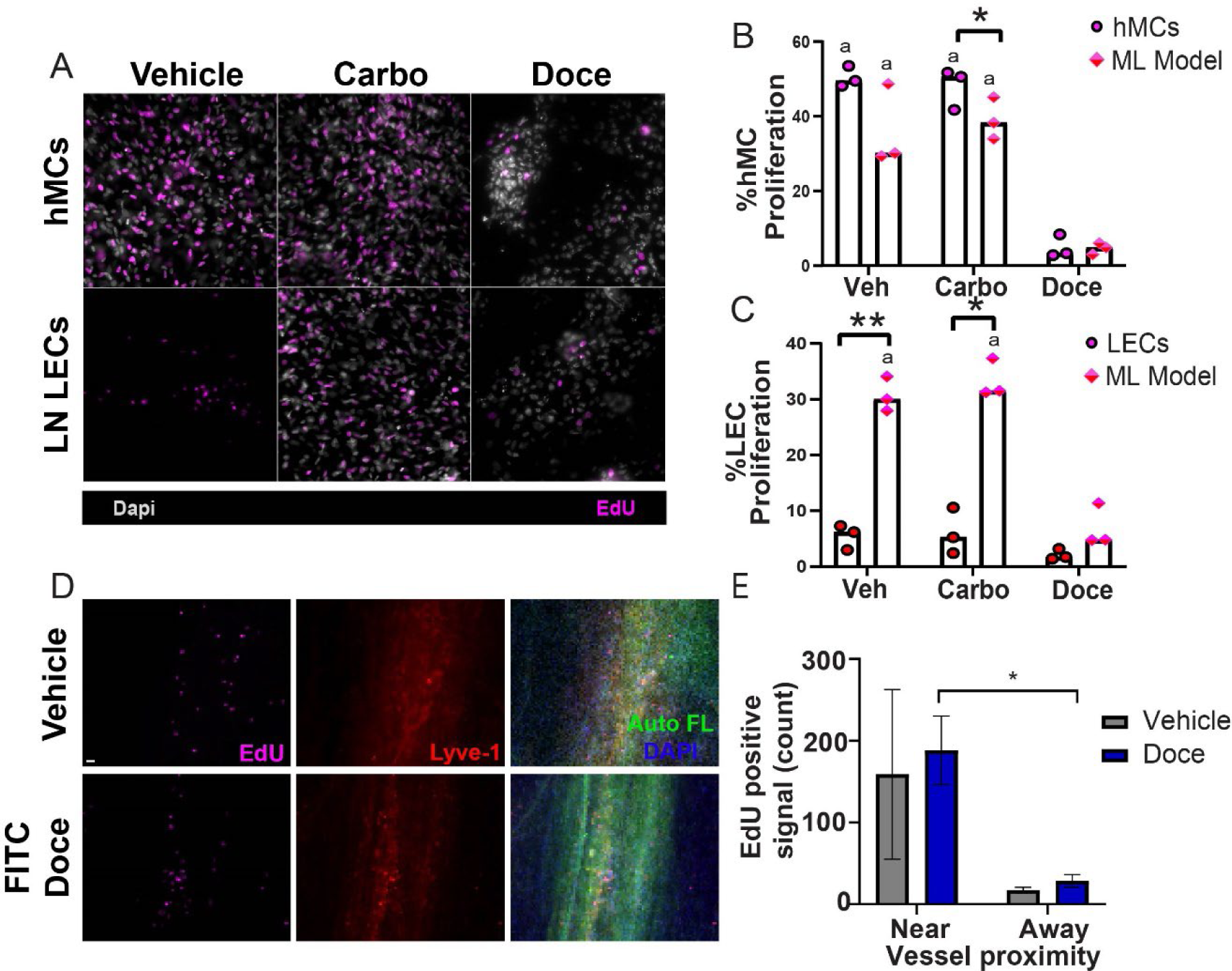
Docetaxel decreases the number of proliferative cells present at 24 hours in vitro with spatial preference in proliferation ex vivo. ML models are treated with 1 µM carboplatin, 1 µM docetaxel, or vehicle (DMSO) for 24 h. During the last 6 h of treatment, 10 µM EdU is added to the culture media. Representative images of proliferation in the ML models are shown, with all nuclei stained with DAPI (gray) and EdU positive cells (magenta). Scale bar is 50 µm (A). Percentage of proliferation of hMCs (B) and LECs (C) is quantified in ImageJ, within the ML model and as monocultures. Data shown are biological replicates (n=3) and mean, with * denoting p<0.05, ** denoting p<0.01. a denotes significant difference from docetaxel treatment. (D) Representative images of proliferation in ex vivo meningeal layers after 18 h culture in vehicle or 1 µM FITC docetaxel and 6 h of treatment of 10 µM EdU is added to the culture media (Scale bar: 50 µm). (E) Quantification of EdU positive cells in meningeal vessels near and away from lymphatic vessels (n=3)

Based on the differences seen in co-culture compared to cell types in isolation in the presence of docetaxel, we wanted to examine if we could capture any spatial effects of docetaxel on proliferation and lymphatic vessel remodeling. To test this, ex vivo meningeal layers were cultured in the presence of 1 µM FITC-labeled docetaxel or DMSO with EdU for 6 h to visualize uptake in the tissue, which was achieved at the start of 3 h (Supplementary Figure S8), prior to a total of 18 h of incubation. Proliferation was not significant between docetaxel and vehicle treatment (Figure 6D,E, Supplementary Figure S9A,B). However, there were spatial differences in proliferation near and away from the vessel, especially in the presence of docetaxel (Figure 6E). Moreover, proliferation was also significantly different between the lymphatic vessels in the transverse sinus and the superior sagittal sinus in the presence of docetaxel, with highest proliferation in the transverse sinus (Supplementary Figure S9C). Although not significant, in the presence of FITC docetaxel from live imaging, remodeling through intussusceptions/loops and branching was lower compared to the vehicle (Supplementary Figure S9D). These remodeling outcomes were different spatially in the transverse sinus compared to the superior sagittal sinus in the presence of the vehicle or FITC docetaxel. In the SSS, branching and loops were not trending differently, whereas these outcomes were lower in the transverse sinus in the presence of docetaxel compared to vehicle.

## Discussion

Although a plethora of tools and experimental methods can be applied to develop a better understanding of the meningeal lymphatics, most have been in vivo studies. We have employed a three-tier model system including in vivo treatments, ex vivo culture of meningeal layers, and an in vitro meningeal lymphatic model. Here, we have performed in vivo systemic chemotherapy treatments in mice before harvesting and examining the meningeal lymphatics on a tissue level. In vivo experiments are ideal for modeling systemic treatment with the physiological complexity of the body, a feature not yet possible in vitro. In vivo study further allows examination of endpoint structural changes and remodeling. Conversely, ex vivo culture of meningeal layers allows for dynamic imaging and uptake in an isolated system that is modeled outside of the body, but still on a tissue level. We are proposing the first ex vivo meningeal lymphatic model in mice.

Finally, there are no existing in vitro models that examine changes in both lymphatic endothelial cells and meningeal fibroblasts together to recapitulate meningeal lymphatic tissue composition. For this purpose, we developed a novel tissue culture insert model of the meningeal lymphatics that allows for examination of immune cell transmigration, cellular crosstalk, cell morphology, and protein secretion. In vitro models are ideal for quick, large screening experiments that allow potential targets to be narrowed down quickly prior to moving into animal work. Each experimental model we have presented provides different levels of complexity and throughput (Table).

For the in vitro model, the goal was to model the lumen of a meningeal lymphatic vessel. The chosen cell types included in this model, MCs and LECs, have specific key functions. Meningeal fibroblasts have a variety of functions ^20^, but generally make up the bulk tissue, maintain meningeal integrity, and contain CSF. Lymphatic endothelial cells are responsible for collecting fluid, altering their junctions and structure to follow the needs of fluid balance ^21^. Culturing LECs in vitro comes with a variety of challenges, and initial attempts to optimize a co-culture media were not as successful as transitioning the MCs to LEC media. Therefore, LEC media was chosen to culture the model. Future applications may require optimization of a different co-culture media, especially in the event of adding additional cells that are also known to be sensitive to media formulation, such as neural cells or immune cells.

|  | In vivo | Ex vivo | In vitro |
| --- | --- | --- | --- |
| Best for | Systemic treatment to mimic patient standard-of-care regimen, endpoint structural changes and remodeling | Dynamic imaging of structural changes and remodeling; uptake, endpoint structural changes and remodeling | Immune cell transmigration, cellular crosstalk, cell morphology, protein secretion |
| Scale | Bulk structure | Bulk structure | Cellular |
| Ease of study | Medium | High | High |
| Ability to test many conditions | Low | Medium | High |
| Complexity | High | Medium | Low |
| Time scale | Fixed time points | Dynamic and fixed | Fixed time points |

Recently, the impact of localized radiation treatment on the meningeal lymphatics has been explored ^22^. However, lymphatic remodeling in response to systemic cancer treatment has not been fully characterized. It is established that platinum and taxane systemic chemotherapy have impacts on ex vivo mesentery lymphatics and lymphatic endothelial cells in vitro ^15,16^. Due to these reports, we wanted to explore the impact of these chemotherapeutic agents on meningeal lymphatic vasculature, especially since the meningeal lymphatics are involved in drainage and cognitive function ^23^. This is notably important since cognitive impairment is commonly reported by patients who have had chemotherapeutic treatment ^24,25^. There is established cognitive dysfunction associated with docetaxel treatment in rodents ^26–28^ and humans ^29,30^. The impact of platinum chemotherapy on cognitive function is less clear ^31,32^.

Here, we show that taxane chemotherapy significantly decreased meningeal lymphatic remodeling in vivo and ex vivo and caused disruption in an in vitro model of the meningeal lymphatics, while platinum chemotherapy did not. In vivo, docetaxel significantly decreased meningeal lymphatic vessel diameter and number of intussusceptions/loops. However, branching points were not significantly different after docetaxel exposure. Ex vivo, docetaxel caused significant decreases in number of intussusceptions/loops and branching points. This could possibly provide insights on fates in remodeling, such as branching still occurring, periendothelial cell networks and splitting into intussusceptions as seen in blood ^33^, and lymphangiogenesis associated with lymphatic development and malfunction being impacted ^34–37^. In contrast, in breast stroma, docetaxel induced lymphatic remodeling when tumor was present but had no impact without the tumor ^15^. Interestingly, though our mouse model had a tumor presence in the body, our in vitro model did not require tumor presence to demonstrate disruption. There is a delicate balance between lymphatic remodeling and function. Docetaxel is known to cause lymphatic dysfunction, with common occurrences of lymphedema following treatment during breast cancer ^38–40^. Prior work has shown that docetaxel accumulates in the lymphatics with both subcutaneous and intravenous delivery, whereas carboplatin does not ^41^. Further, docetaxel cannot cross the blood-brain barrier (BBB), suggesting it could accumulate in the meninges and then interact at high concentrations with the meningeal lymphatics. Though docetaxel cannot cross the BBB in its normal form, it potentially can disrupt the BBB, as docetaxel is associated with higher occurrence of breast cancer metastasis to the brain ^42^. Thus, these findings suggest that docetaxel may have uniquely deleterious impacts on meningeal lymphatics.

We further examined the impact of platinum chemotherapy on meningeal lymphatics. Previously, we reported that carboplatin induced remodeling in lymphatics in healthy non-cancerous rat mesentery, mouse mammary fat pads, mouse lungs, and human omentum ^16^. Contrasting our previous findings, platinum therapy did not induce increased meningeal lymphatic remodeling compared to vehicle in vivo or ex vivo, or cause disruption in our in vitro model. This could be due to the location of lymphatics and function determining responses to treatment and outcomes. For instance, lymphatics in different parts of the body are represented in various environments, chemokines, and flow and drainage conditions. Thus, different responses to perturbations could potentially cause a difference in the meningeal lymphatics specifically. Also, the proximity to where the therapy is administered could possibly have an increased effect on remodeling outcomes. There are four options for treatment location in central nervous system cancers, including intrathecal CSF administration, intraventricular therapy, localized, and regional therapy. Future studies could explore how different treatment locations impact meningeal lymphatics.

Further, there is potential for differences in LEC response based on source. Within our in vitro model, we used lymph node LECs, specialized cells that are highly relevant in our studies, but other types, such as dermal LECs, could respond differently based on their physiological function. In our studies, carboplatin disrupts LN LEC junctions but does not increase proliferation. Docetaxel disrupts junctions and decreases proliferation. We previously reported that dermal LECs are activated by carboplatin and demonstrate disrupted junctions, but that docetaxel does not disrupt junctions in dermal LECs^16^. Prior reports show that dermal LECs have a decreased ability to migrate and form tubes when treated with docetaxel ^43^. Thus, LEC source may impact response to chemotherapy.

In our ex vivo model, we observed that the signal was weaker in both Prox1-Tdtomato mice as well as LYVE-1 staining in ex vivo studies compared to in vivo studies. LYVE-1 is a receptor for hyaluronic acid (HA) and the signal could have been impacted by being outside of the environment without access to HA. Still, the ex vivo model showed more sensitivity in changes to branching in the presence of chemotherapeutic agents than seen with in vivo.

We also observed differences in remodeling outcomes based on location of the lymphatic vessels. With the transverse sinus and superior sagittal sinus, recent studies noted differences of remodeling and functionality at the two different locations ^19,44–46^. There were no significant differences between the average number of branches and loops in terms of remodeling in both locations along the meningeal layer at the superior sagittal sinus. However, there was a significant decrease of loops at the transverse sinus. Waste and fluid flow, which contains the chemotherapy, drains through the meningeal lymphatics in the CNS with the superior sagittal sinus ultimately draining to the transverse sinus^47^, so the level of remodeling could possibly correlate with the drainage location and be further explored with tracers in the presence of chemotherapeutic agents in future studies.

We further utilized our in vitro and ex vivo models to examine how docetaxel and carboplatin impact proliferation in the meningeal lymphatics. Carboplatin, a platinum-based chemotherapy, prevents proliferation by binding to DNA and inhibiting replication and triggering apoptosis ^48^. Docetaxel, a taxane chemotherapy, interferes with microtubules, leading to cell cycle arrest and preventing cell division ^49,50^. These chemotherapies are known to have off-target side effects, as they can often impact rapidly dividing healthy cells, such as hair, nails, and skin. Specifically, fibroblasts are highly proliferative cells that are prevalent all over the body, but also in the bulk structure of the meninges and surrounding meningeal lymphatic vessels. Thus, they could also be impacted during treatment based on their proliferative nature. Our in vitro model demonstrated a decrease in percentage of proliferating meningeal cells in docetaxel-treated groups compared to both the vehicle and the carboplatin, both alone and in the meningeal lymphatic model.

In contrast, LECs only exhibited decreased proliferation when treated with docetaxel and in co-culture with meningeal cells. It is interesting to note that we previously reported that dermal LECs are activated by carboplatin and demonstrate a threefold increase in proliferation (Harris). This demonstrates the importance of LEC source and microenvironment in altering response to chemotherapy. Interestingly, the ex vivo model did not demonstrate differences in proliferation during docetaxel treatment but did exhibit a spatial preference for proliferation near lymphatic vessels, suggesting important crosstalk between LECs and meningeal cells. Overall, we have developed in vitro and ex vivo models that further validated our in vivo findings, demonstrating that docetaxel has negative impacts on the meningeal lymphatics.

Looking forward, the in vitro meningeal lymphatic model presented within can be tailored to a wide variety of applications, such as modeling different pathological conditions with inflammation or tumor cells or altering the orientation of the cells to model afferent vs efferent vessels. This model can also be combined with photo-crosslinkable hydrogels to represent the brain parenchyma. Additionally, the meninges are an immune cell reservoir, so incorporating immune cells such as macrophages and lymphocytes is of great interest toward increasing complexity. This model also could employ specific lymphatic endothelial cells and fibroblasts from different locations in specific parts of the body to assess effects of treatment and/or perturbations from a localized area. Future applications of our ex vivo model could be enhanced by perfusion with a bioreactor to better represent the natural lymphatic system with fluid flow. This would allow assessment of the impact of chemotherapeutics and fluid flow response on remodeling outcomes to simulate physiological conditions.

## Conclusion

We have presented a three-tiered approach toward studying the meningeal lymphatics, which is an area of great interest in understanding CNS malignancy. In vivo studies allow for systemic treatment and tissue-level analysis. The ex vivo component allows for dynamic imaging and higher ease of experimentation while also providing tissue-level insight. Our novel in vitro meningeal lymphatic model allows examination on a cellular level, higher throughput, and customization based on application. Further, we have shown that the meningeal lymphatics have differing responses to platinum and taxane chemotherapy compared to other lymphatic systems in the body, thus demonstrating off-target impacts on the CNS that require future study.

## Supporting information

Z-stack capture of optimal plane for analysis

## Author Contributions

L.M.R., J.H.H., M.R. and J.M contributed to the conceptualization, methodology, and preparation, and editing of the manuscript. F.A., T.F., and M.R. conducted in vivo tumor implantation and chemotherapeutic treatment. L.M.R., F.A., T.F., and M.R. harvested tissues from in vivo studies. L.M.R. performed meningeal layer dissection, tissue processing, staining, and imaging for in vivo and ex vivo studies and analysis. J.H.H. developed the in vitro tissue engineered model and performed staining, imaging, and analysis. J.J.C. developed the Cellprofiler pipeline for diameter quantification for meningeal layers.

## Acknowledgements

NIH National Institute of Aging grant AG071661, CCR grant from Susan G Komen CCR17483602 and 1R01CA253285 and 1R01CA262634 from the NCI, with aid from Grant #IRG 81-001-26 from the American Cancer Society. TF was supported by a trainee fellowship from the University of Virginia Comprehensive Cancer Center. JH was supported by the Virginia Tech Institute for Critical and Applied Science (ICTAS).

