## Supplemental Figures and Information for "Demonstration of chemotherapeutic mediated lymphatic changes in meningeal lymphatics in vitro, ex vivo, and in vivo"

### Supplementary Methods

*Viability Assays*

LECs are seeded in fibronectin coated 24 well plates and cultured in various experimental media. After 24 hours, viability was quantified via LIVE/DEAD kit, as specified in main methods.

*Image Quantification*

Using Fiji, LECs were traced, and area and aspect ratio were recorded. For total area of fluorescence, images were thresholded using Fiji and total area of signal was measured. Cells with disrupted junctions were counted and reported as a percentage of total cell count.

### Supplementary Figures

**Supplementary Video S1.** Z-stack capture of optimal plane for analysis. Representative z-stack of meningeal layer labelled for CD31 (cyan), Lyve-1 (red), α-SMA (green), and DAPI (blue).

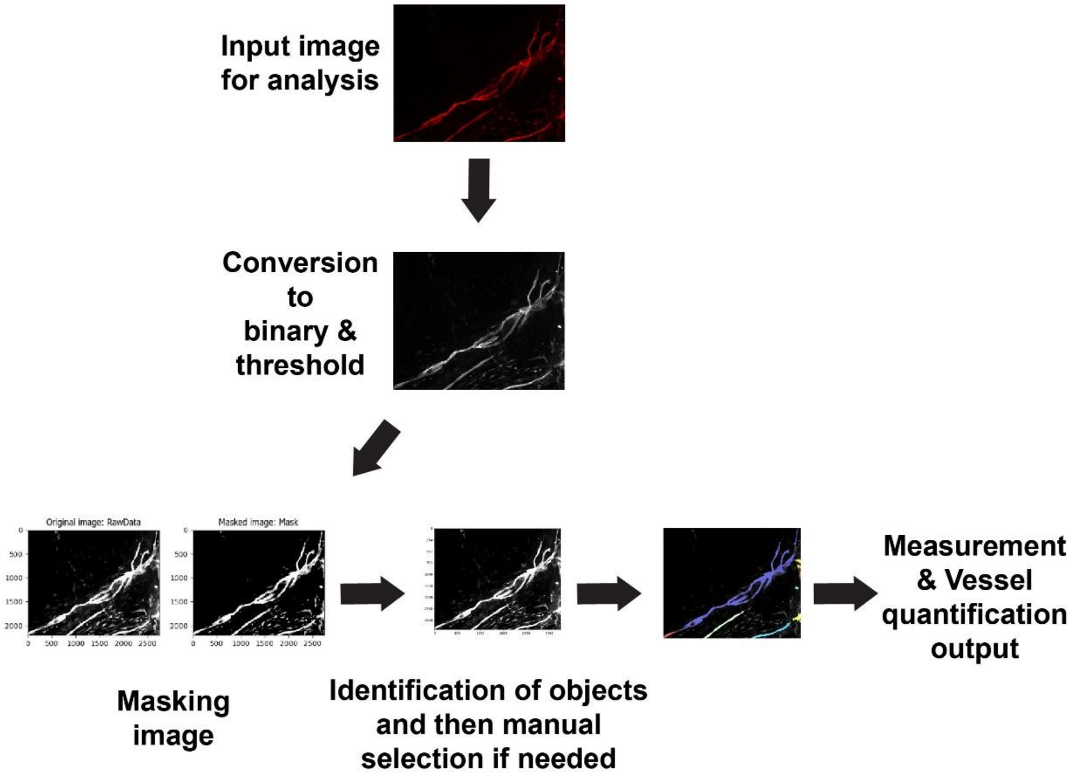


**Supplementary Figure S2. Workflow for automated image analysis through Cellprofiler pipeline.** Schematic on Cellprofiler pipeline outlining modules used for quantification of meningeal layer vessel diameters.

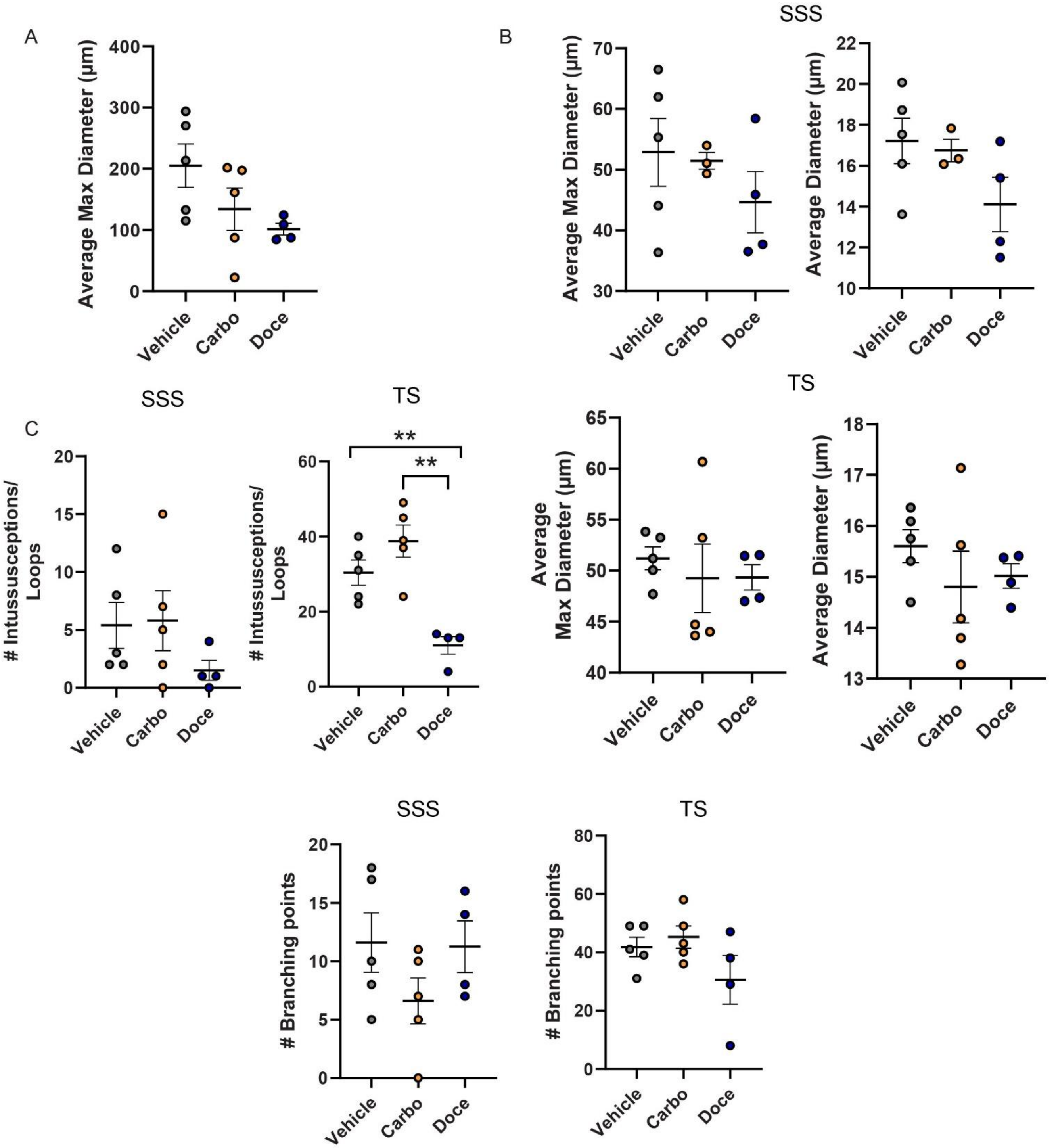


**Supplementary Figure S3. Remodeling through intussusceptions is different across chemotherapeutics in the TS.** (A) Quantification of the maximum diameter in the presence of 1 µM carboplatin, 1 µM docetaxel, or vehicle (DMSO) and average max diameter. (B) Quantification of the maximum diameter in the presence of 1 µM carboplatin, 1 µM docetaxel, or vehicle (DMSO) and average max and average overall diameter. (C) Quantification of the maximum diameter in the presence of 1 µM carboplatin, 1 µM docetaxel, or vehicle (DMSO) and average max diameter in the SSS or the TS location in the meningeal layer. (D) Quantification of intussusceptions/loops and branching in the presence of 1 µM carboplatin, 1 µM docetaxel, or vehicle (DMSO) in the SSS or TS location in the meningeal layer. All graphs represent Mean ± SEM (n=5 for vehicle and carboplatin; n=4 for docetaxel, **, $p < 0.01$).

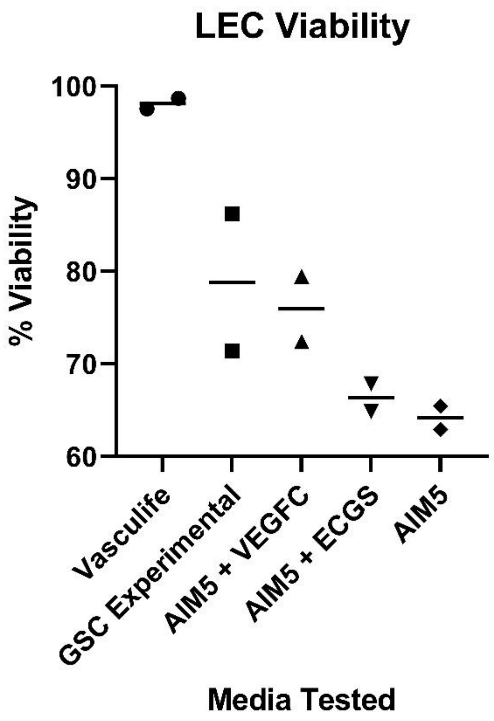


**Supplementary Figure S4. LECs are not viable in other tested experimental medias.** Viability of LN LECs after 24 h in various cell culture medias, as determined by LIVE/DEAD kit. n=2.

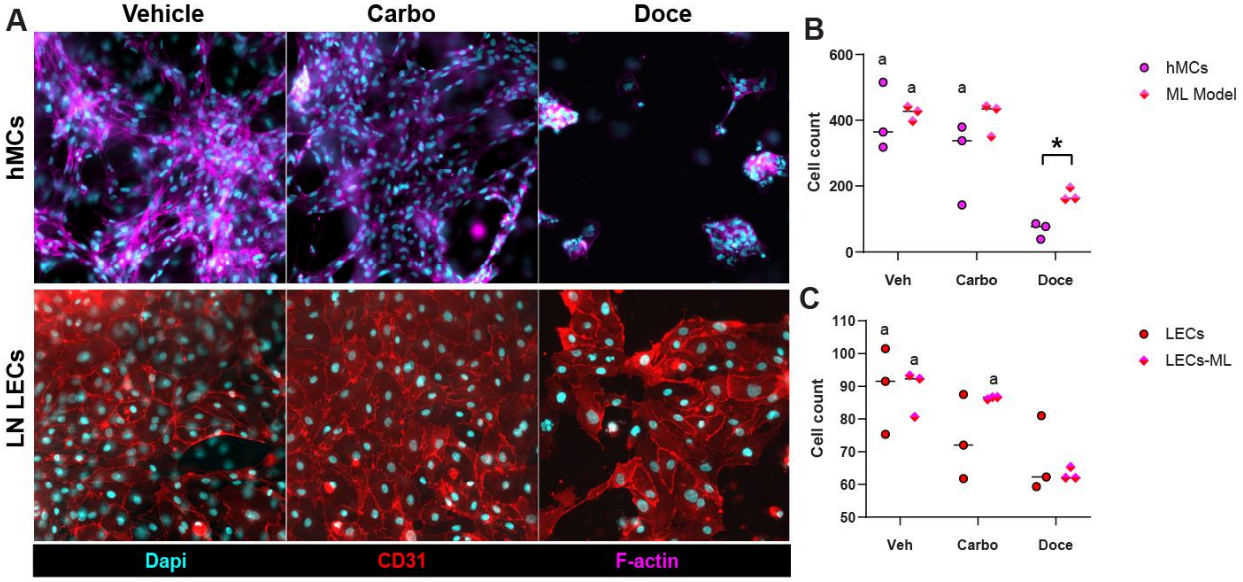


**Supplementary Figure S5.** ML models are treated with 1 µM carboplatin, 1 µM docetaxel, or vehicle (DMSO) for 24 h. Representative images of monocultures controls are shown, with nuclei stained with DAPI (cyan), hMCs stained for F-actin (magenta) and LECs stained for CD31 (red). Scale bar is 50 µm (A). Cell count of hMcs (B) and LECs (C) within the ML model and as monocultures after treatment. Data shown are biological replicates (n=3) and mean, with * denoting p<0.05, ** denoting p<0.01. a denotes significant difference from docetaxel treatment.

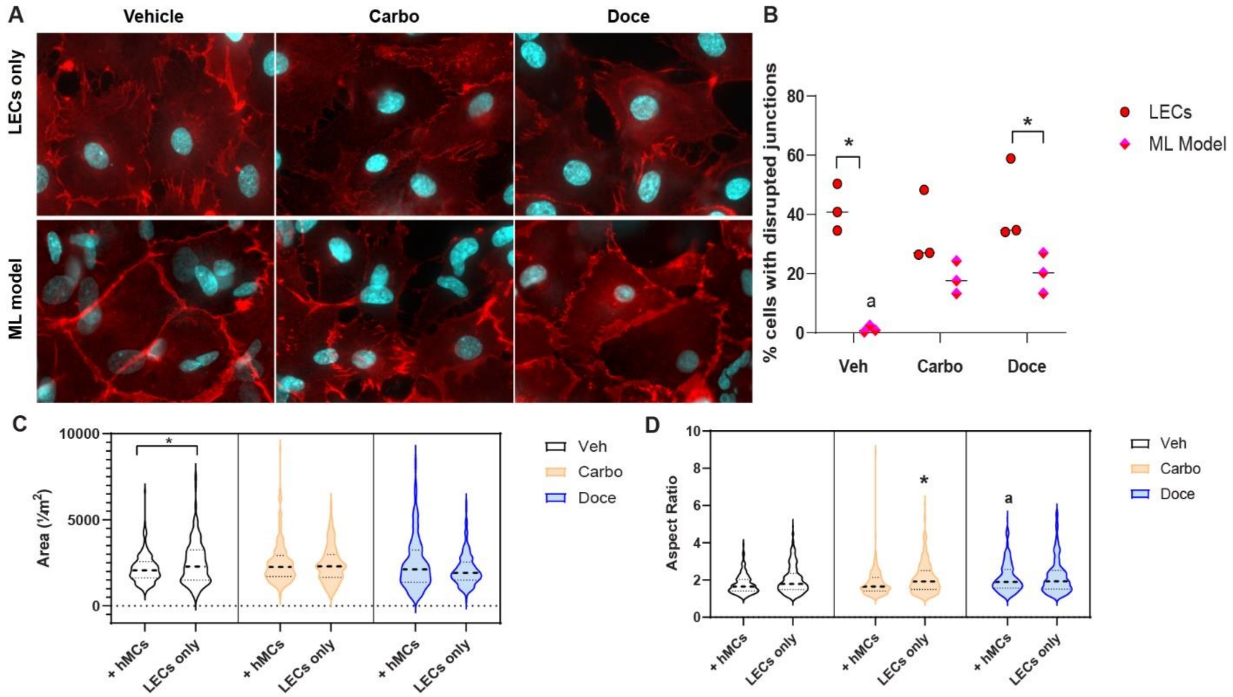


**Supplemental Figure S6. LEC junctions and morphology are impacted by chemotherapy treatment and co-culture.** ML models are treated with 1 µM carboplatin, 1 µM docetaxel, or vehicle (DMSO) for 24 hours. Representative images of LECs are shown, with nuclei stained with Dapi (gray) and LECs stained with CD31 (red). Percentage of LECs with disrupted junctions is quantified (B), as well as LEC area (C) and aspect ratio (D). Data shown are biological replicates (n=3) and mean for disrupted junctions. For LEC morphology, 80 LECs from each biological replicate are quantified and displayed, but statistical analysis was performed with the average. For statistics, * denotes p<0.05 and a denotes significant difference from docetaxel treatment.

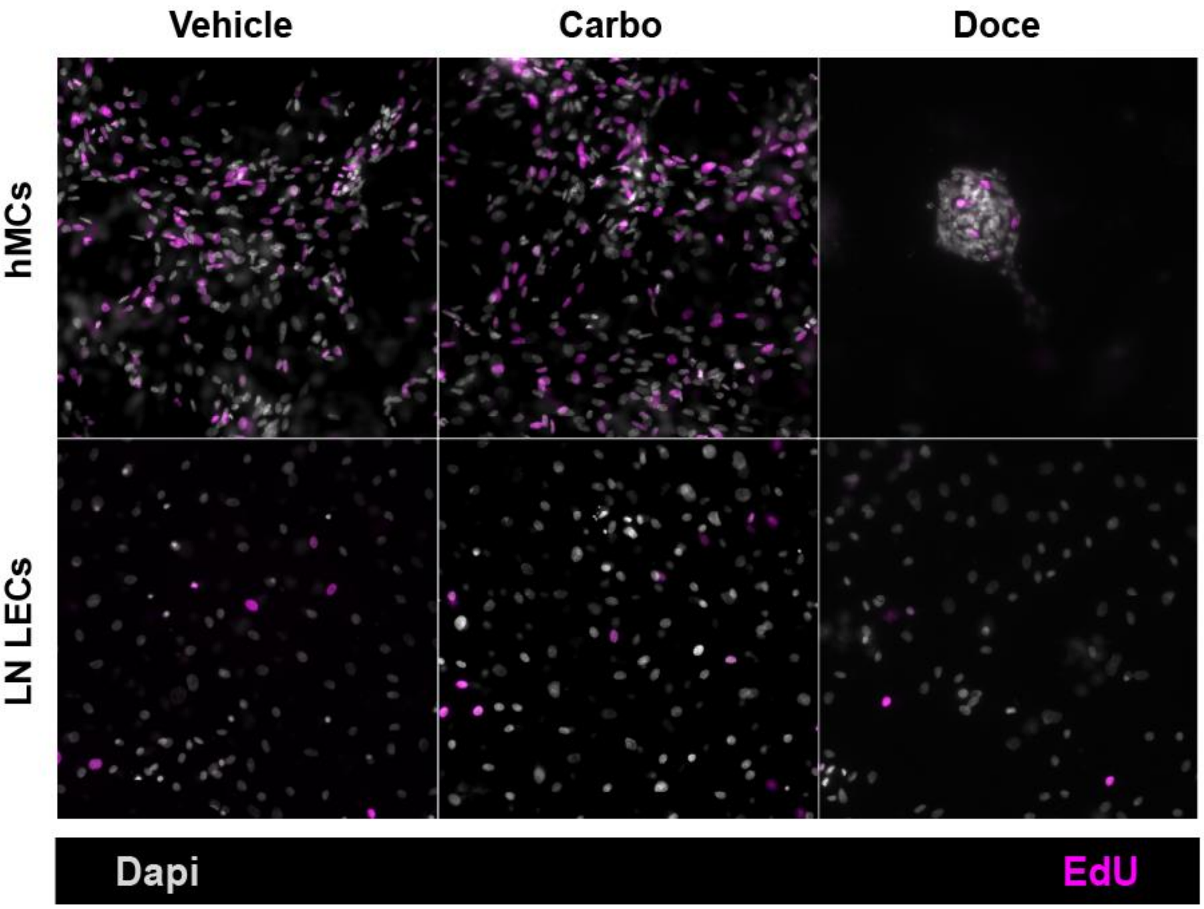


**Supplemental Figure S7. Proliferation in monoculture controls.** ML models are treated with 1 µM carboplatin, 1 µM docetaxel, or vehicle (DMSO) for 24 hours. During the last 6 h of treatment, 10 µM EdU is added to the culture media. Representative images of proliferation in the monoculture controls are shown, with all nuclei stained with Dapi (gray) and EdU positive cells in magenta. Scale bar is 50 µm (A).

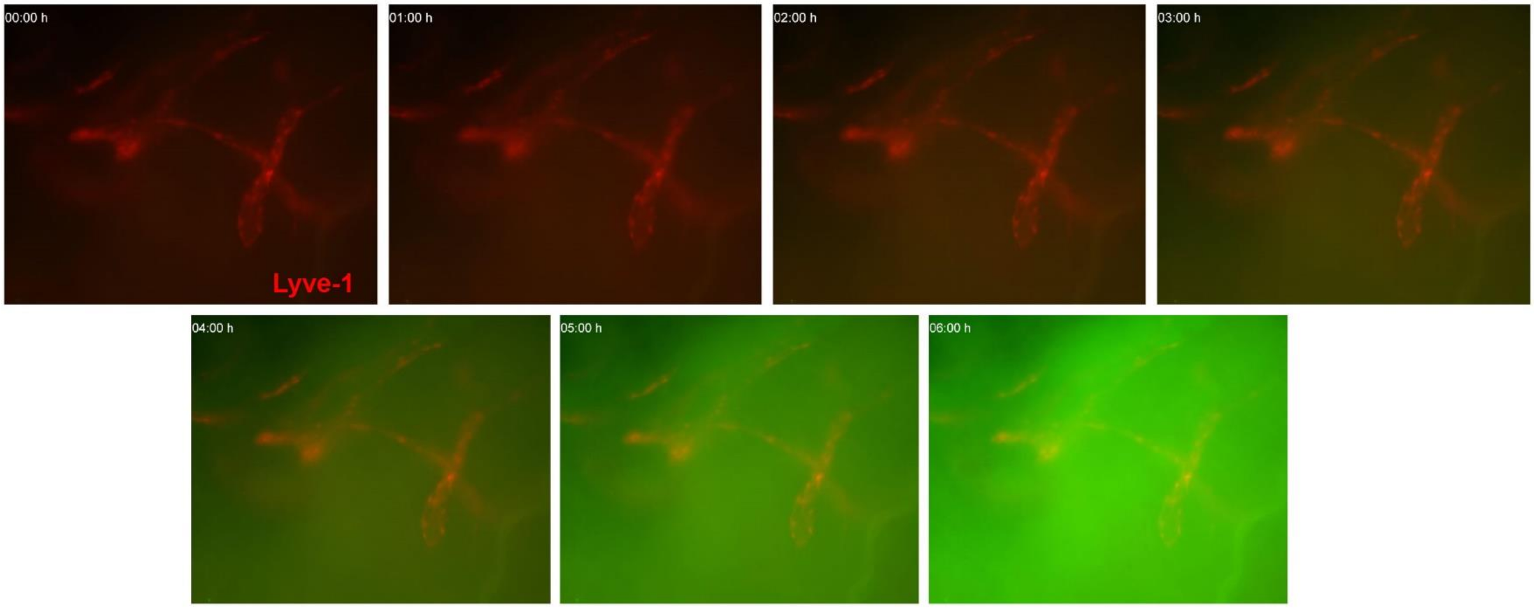


**Supplementary Figure S8. Uptake in the presence of docetaxel in an ex vivo meningeal layer begins at 3 h.** Snapshots of time points after the meningeal layer (Lyve-1 staining in red and 1 µM FITC docetaxel in green) was added to the plate during live imaging and allowed to incubate over the course of 6 h (n =3).

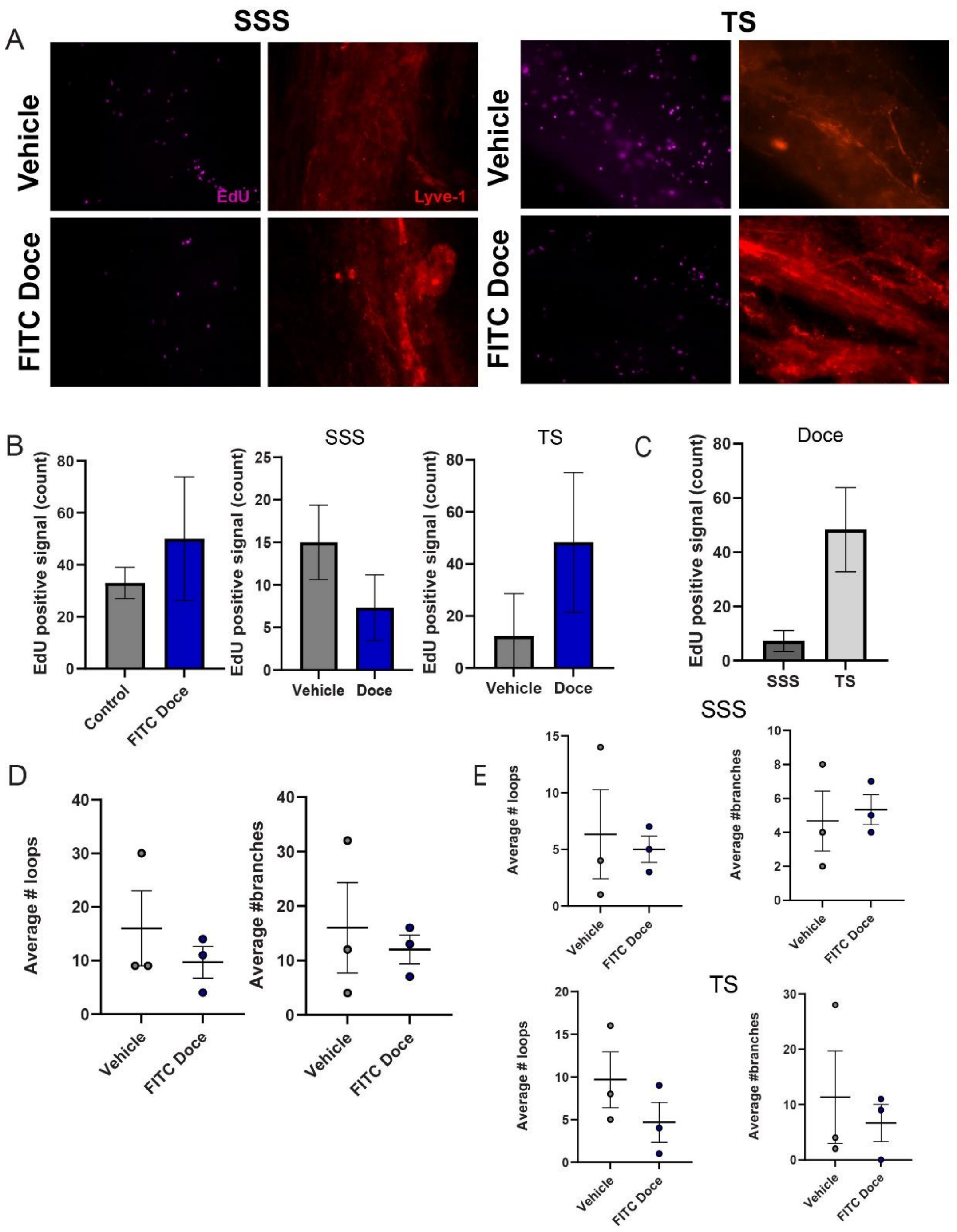


**Supplementary Figure S9. Proliferation and remodeling reveal a decreasing trend in the presence of docetaxel ex vivo.** (A) Representative images of EdU and Lyve-1 in ex vivo meningeal layers in the presence of vehicle or 1 µM FITC docetaxel in the SSS or the TS. (B) Quantification of total EdU positive signal in lymphatic vessels between vehicle and FITC Docetaxel (left) and differences in the vessels in the SSS or TS. (C) Quantification of total EdU positive signal in the FITC-docetaxel treated meningeal layers in the SSS and TS. (D) Quantification of the average intussusceptions and branching points in vehicle or FITC docetaxel–treated layers. (E) Quantification of remodeling outcomes based on spatial location of SSS or TS. All graphs represent Mean ± SEM.
